# Vestibular involvement in transcranial electrical stimulation: body sway as a marker of unintended stimulation

**DOI:** 10.1101/2024.09.13.612582

**Authors:** Ilkka Laakso, Janita Nissi, Otto Kangasmaa

## Abstract

**Background:** Transcranial electrical stimulation (tES) techniques are widely used to modulate brain excitability, though their mechanisms remain unclear.

**Objective:** We aimed to demonstrate that an alternating current applied to several commonly used tES electrode montages can influence the vestibular system.

**Methods:** Body-sway responses during alternating current stimulation at 4.6 or 4.8 Hz were measured in eight participants standing on a force platform. Sham and five active conditions, including common motor cortical tES electrode montages, were applied in a double-blind experiment.

**Results:** All active conditions increased lateral body sway compared with sham at the stimulus frequency, with very large to huge effect sizes. The sway magnitude was proportional to the montage-specific electric field within the vestibular system, obtained from a computational model. The frequency and phase of the body sway were locked to the stimulus frequency and phase. An additional experiment in three participants showed that the sway response was the largest in the lower legs, similarly for both tES with a motor cortical electrode montage and an electrical vestibular stimulation montage. Together, the results suggest a direct effect of tES on the vestibular system.

**Conclusion:** Unwanted vestibular effects may interfere with the interpretation of tES studies across a wide range of typical tES frequencies and current intensities.

**Highlights:**

- Alternating current stimulation produces a lateral body sway at the stimulation frequency.
- Larger lateral electric field in the vestibular system produces a larger sway.
- Several common tES montages can directly influence the vestibular system.
- Unwanted vestibular effects may complicate the interpretation of tES studies.

## Introduction

Transcranial electrical stimulation (tES) including its direct (tDCS) and alternating current (tACS) variants are popular techniques for studying and modulating the excitability of the brain. These techniques are applied in both research and clinical settings to influence neural activity, yet their mechanisms are still unclear.

Interestingly, tES shares similarities with electrical vestibular stimulation (EVS), another non-invasive electrical stimulation technique [1, 2], such as the use of similar electrodes and currents of comparable frequency and magnitude. Despite these similarities, tES is generally thought to affect the brain, while EVS is known to directly influence the vestibular system, inducing postural changes and eye movements [2]. For example, direct current (DC) applied through electrodes attached behind the ears makes the subject lean in one direction, while an alternating current (AC) makes them sway from side to side. These responses are easily observable using a force platform or by analysing movements from a video of the subject.

EVS typically uses current intensities in the range of ≥ 1 mA [2], but a few hundred µA may be sufficient to produce postural sway [3]. Similarly, tES commonly uses 1–2 mA current intensities [4]. Even though tES electrodes are not typically placed near the inner ear, the current could still spread through the head tissues to the vestibular organ, potentially generating vestibular responses. Indeed, computational modelling suggests that 2 mA current applied with the C3–Fp2 montage, commonly used for stimulation of the somatomotor cortices, produces an average electric field strength of 0.20 ± 0.02 V/m over the bilateral vestibular systems (Supplementary Fig. 1b). For comparison, 1 mA current applied with a typical bilateral EVS montage would generate a very similar field strength of 0.18 ± 0.03 V/m (Supplementary Fig. 1b).

Based on these modelling data, we hypothesized that tES with various electrode montages could directly affect the vestibular system, leading to an EVS-like body sway. Furthermore, we expected that the sway magnitude would be proportional to the electric field within the vestibular system so that stronger fields would lead to a larger sway. To test these hypotheses, we measured body sway responses in eight participants in a pre-registered (https://doi.org/10.17605/OSF.IO/W7EZX) double-blind experiment.

## Methods

The present study consisted of two experiments and computational modelling.

### Participants

Eight participants (3 male, 5 female, 19–29 years) took part in experiment 1, and three participants (2 male, 1 female, 28–41 years) took part in experiment 2.

All participants were in good general health, reported no history of vestibular dysfunction and had no contraindications to transcranial stimulation. Each participant gave their written informed consent and received a lunch ticket valued less than 20 euros for their participation. Ethical approval for the study was obtained from Aalto University Research Ethics Committee (diary number D/1219/03.04/2022). All procedures were conducted in accordance with the Declaration of Helsinki.

### Transcranial electrical stimulation and force platform measurements

In both experiments, the stimuli were AC sine waves lasting 120 s with a ramp-up and ramp-down (5 s in experiment 1 and 3 s in experiment 2). The stimuli were applied using a Starstim 20 stimulator (Neuroelectrics, Barcelona, Spain) with 25 cm^2^ round saline-soaked electrodes placed at various locations on the scalp. All reported current intensities are peak values.

For recording the sway response, the participant stood on a 1.5 cm thick plastic foam mattress placed on a force platform (Wii Balance Board, Nintendo, Japan) with eyes closed, arms crossed, shoes off, and feet together. The balance board was modified by soldering additional wires to the circuit board so that four raw amplified pressure sensor signals could be directly recorded. An eight-channel oscilloscope (PicoScope 4824A, Pico Technology Ltd., UK) recorded the sensor voltage signals at a sample rate of 1 kHz with 12 bit precision, range of ±100 mV, and AC coupling. The stimulation current waveforms were synchronously recorded in a non-contact way using current transformers (CT-C0.1-BNC, Magnelab Inc., CO, US).

The voltage signals from the four pressure sensors were converted to a centre-of-pressure (CoP) signals in the lateral and anterior-posterior (AP) directions by addition and subtraction, and scaling them with the lateral and AP sensor distances (0.430 m and 0.235 m). As we used the raw sensor voltages without calibration, we reported the results in arbitrary units.

### Experiment 1: Does tES increase body sway compared with sham?

This experiment was double-blind and counterbalanced, and the methods and analyses were pre-registered prior to data collection (https://doi.org/10.17605/OSF.IO/W7EZX). Power calculations using effect size estimates (Cohen’s *d*: 1.17–1.8) of experiment 2 showed that four to seven participants should give ≥ 80% power for finding an increased sway compared with sham (one-tailed tests) with 0.05 significance level for all studied electrode configurations, so the sample size was selected as eight.

We expected that the body sway response should appear at any suitable frequency, so instead of fixing the stimulation frequency a priori, an automated MATLAB (The Mathworks, Inc.) script generated the stimulus waveforms for two randomly selected frequencies from among 4.2, 4.4, 4.6, 4.8, and 5.0 Hz such that both the participants and the researchers were blinded to the frequency until the data collection was finished. The realized frequencies were *f*_1_ = 4.6 Hz and *f*_2_ = 4.8 Hz.

Four electrodes were placed on the scalp at C3, C4, Fp2 of the 10-20 system, and over the left cerebellar hemisphere 4 cm lateral from the inion. In the following, the latter electrode location is denoted ‘I1’ according to the closest electrode of the 10-5 system [5]. AC stimulus at *f*_1_ or *f*_2_ was applied to various pairs of electrodes such that there were six conditions: I1-Fp2 with 1.5 mA (*f*_1_), I1-C4 with 1.5 mA (*f*_1_), C3-Fp2 with 2.0 mA (*f*_1_), C3-C4 with 2.0 mA (*f*_1_), C3-Fp2^*†*^ with 2.0 mA (*f*_2_), and sham. For sham stimulus (applied to C3-Fp2), the stimulus was ramped during 5 s up to 2.0 mA and then immediately ramped down during the following 5 s. The C3-Fp2^*†*^ condition was otherwise the same as C3-Fp2 but used a different stimulation frequency. The order of the conditions was counterbalanced using an 8×6 Latin rectangle and randomized using an automated Python program.

Prior to the start of the measurements but after attaching the electrodes, while the participant was still seated, 10 s (with 3 s ramp up and down) of 1.5 mA AC at 4 Hz was fed through each electrode montage to verify the electrode impedances and to familiarize the participant to stimulation. An additional unblinded control condition, lasting approximately 130 s, with no stimulation was applied before the other conditions to allow the participant to get used to the measurement setup.

From the CoP signals (130 s), we trimmed away the initial 15 s and the last 5 s. For the CoP data of the control condition, we cut away data from the beginning and end of the recording so that the remaining duration was the same as that for the other CoP signals. From each trimmed signal, we then calculated the power spectrum (`pspectrum` in MATLAB) with 0.1 Hz frequency resolution and obtained the value of the spectrum at stimulation frequency.

Statistical tests were preregistered prior to the data collection, and MATLAB was used for all analyses. The level of statistical significance was chosen as 0.05, and p-values were reported raw, without corrections for multiple comparisons. Generalized extreme Studentized deviate test for outliers were used, and if outliers were found, tests were reported both with and without outliers. We conducted the following confirmatory analyses.

To confirm whether active conditions increased body sway compared with sham, the log-transformed sway magnitude at the stimulation frequency of each active condition was compared to that of sham using paired t-tests (one tailed). Sham data was tested against the control data using a paired t-test (two tailed). The process was repeated for the spectral components calculated for the lateral and AP sway signals.

To confirm the effect of the calculated vestibular electric field strength on the sway magnitude, we planned to fit a linear mixed effects model with specification `“logSway ∼ 1 + EF + (1+EF| Subject)”` to the data, where `logSway` is the log-transformed spectral component value at the stimulation frequency, and `EF` is the mean of the electric field magnitude over both vestibular systems for each active montage (continuous variable). We originally planned to use a preregistered set of electric field values, but after noticing a small error in the original calculations, we instead used newly calculated values that slightly differed from the preregistered values (1.9% relative rms difference). Our preregistered plan was that, if the full model failed to converge, we would change the covariance pattern to diagonal. If the model still failed to converge, the random effects structure would be changed to an intercepts-only model. The former alternative plan realized for both lateral and AP sway signals, and we thus used the diagonal covariance pattern for both models. The effect of the electric field magnitude was tested using a likelihood ratio test comparing the full model to an alternative model that lacked the fixed effect of the electric field.

We performed the following exploratory analyses that were not included in the preregistration. To explore whether the frequency of the sway differed from the frequency of the stimulus, we used the `findpeaks` function in MATLAB to find the largest peak in range 4.8 ± 0.5 Hz from the power spectrum of the lateral sway signal. We then calculated the mean and standard deviation of the difference between the frequency of the largest peak and the stimulation frequency.

To explore the differences between the phases of the stimulus and response, we calculated the phase shift of the lateral sway signal from the stimulation waveform using discrete Fourier transforms at the stimulation frequency. To confirm whether the phase shift was constant between all active conditions within each participant, we calculated the standard deviation of the error from a participant-specific constant model.

As the confirmatory analyses showed that the mean of the electric field magnitude was an invalid model for the sway magnitude, we explored which alternative measures of the electric field were better predictors of the sway magnitude, using the same linear mixed effect model analysis as described above but replacing `EF` with an alternative measure.

### Experiment 2: Which body segments are affected by tES and EVS?

This experiment aimed to investigate how the body sway signal is distributed among body segments and compare the sway produced by tES to that produced by EVS. The experiment was performed prior to experiment 1 but is reported last for presentational purposes. The experiment was unblinded, but reporting the results was still deemed appropriate, as the results of experiment 1 showed extreme effect sizes. Furthermore, the outcome (body sway locked exactly to the stimulation frequency) would be difficult if not impossible to produce through any known wilful or unintentional processes except as a direct effect of stimulation.

An AC stimulus at 4 Hz was applied in six conditions in the following order: no stimulus as a negative control, tES with 1.0, 1.5, and 2.0 mA given through electrodes at C3 and Fp2 of the 10-20 system; another negative control; and EVS as a positive control with 1.0 mA applied through electrodes attached to the bilateral mastoid processes.

During stimulation, the participant stood on the force platform facing a smartphone (Samsung Galaxy s23+) fixed to a tripod. The smartphone filmed the participant’s movements with 1920×824 resolution at 120 frames per second (Android 14 Camera). The experiment was performed in a green room to aid video analysis.

The participant’s silhouette was segmented from the video data using MATLAB. The body was evenly divided into 12 regions of interest (ROI) along the inferior-superior axis, and the horizontal centre of gravity (CoG) of the pixels of the silhouette was calculated in each ROI. Finally, we calculated the power spectrum of the CoG signal over 110 s, trimming away the start and end of the video.

As the purpose of the experiment was illustrative with only three participants, no statistical analyses were performed. However, the effect size estimates were used for selecting the sample size for experiment 1.

### Computational modelling

Computational modelling was used to estimate the electric field in the vestibular systems for each electrode configuration and to select the configurations for the experiments as well as for analysing the results.

The methods for the electric field calculations were identical to those described earlier [6, 3], but we used 15 models of the head (22–40 years, 11 male, 4 female) that were created based on magnetic resonance imaging (MRI) data obtained with a 3 T MRI scanner (Magnetom Skyra; Siemens Ltd., Erlangen, Germany; Aalto University Research Ethics Committee, diary no. D/574/03.04/2022). The finite-element method with 0.5 mm cubical first-order elements was used to calculate the electric field within the head [7]. The surroundings of the inner ear were modelled with 100 µm cubical elements, such that the current density obtained from the whole-head calculations with the 0.5 mm resolution was used as the Neumann boundary condition for the fine-resolution calculations. The tissue conductivity values were (unit: S/m) [6]: blood 0.7, cancellous bone 0.027, cortical bone 0.008, dura mater 0.35, grey matter 0.2, white matter 0.14, cartilage 0.18, cerebrospinal fluid 1.8, fat 0.08, muscle 0.35, and scalp 0.4.

The inner ear model was created by first roughly segmenting the inner ear from the T2-weighted MRI data and then fitting a template model obtained from the MIDA dataset [8]. The fitted template was used to improve the inner ear model, which was further segmented to the vestibulocochlear nerve, cochlea, and vestibular system (Supplementary Fig. 1a).

As there was no information of the electrical properties of the tissues and fluids within the vestibular system, we used the Maxwell-Garnett formula to assign macroscopic electrical conductivity of the inner ear as a random mixture of spherical CSF inclusions in a cortical bone medium. The volume fraction of the inclusions was chosen to be 0.5, as the volume of the inner ear model (Supplementary Fig. 1a) likely overestimated the real volume. Therefore, the vestibular electric field reported herein is a macroscopic field in a homogenized medium. The field affecting the tissues would be the local microscopic field, which is the sum of the macroscopic field and a secondary field generated by local material inhomogeneities. The microscopic field is linearly proportional to the macroscopic field, which is thus a useful surrogate for potential microscopic effects within the vestibular system.

For reporting, we calculated the average of the electric field magnitude or each electric field component over the entire left or right vestibular system, i.e., the cochlea and the vestibulocochlear nerve were excluded from the average. The head models were rotated such that the directions of the lateral, AP, and superior-inferior axes matched those of the MNI space. The modelling results for five electrode montages considered in this study are reported in Supplementary Fig. 1. When analysing the effect of the electric field on the body sway using linear mixed effect models, we used the average values calculated over the 15 models, i.e., each experimental participant received the same electric field value, ignoring potential inter-individual differences. All reported electric fields are peak values.

## Results

### Experiment 1

Sinusoidal AC was applied through four electrode montages (Fig. 1b) while the participants were standing on a force platform (Fig. 1a). The stimulation produced a clear peak visible in the spectrum of the lateral sway signal at the stimulation frequency, 4.6 or 4.8 Hz, easily observable from the spectrum (Fig. 1c).

**Figure 1.**
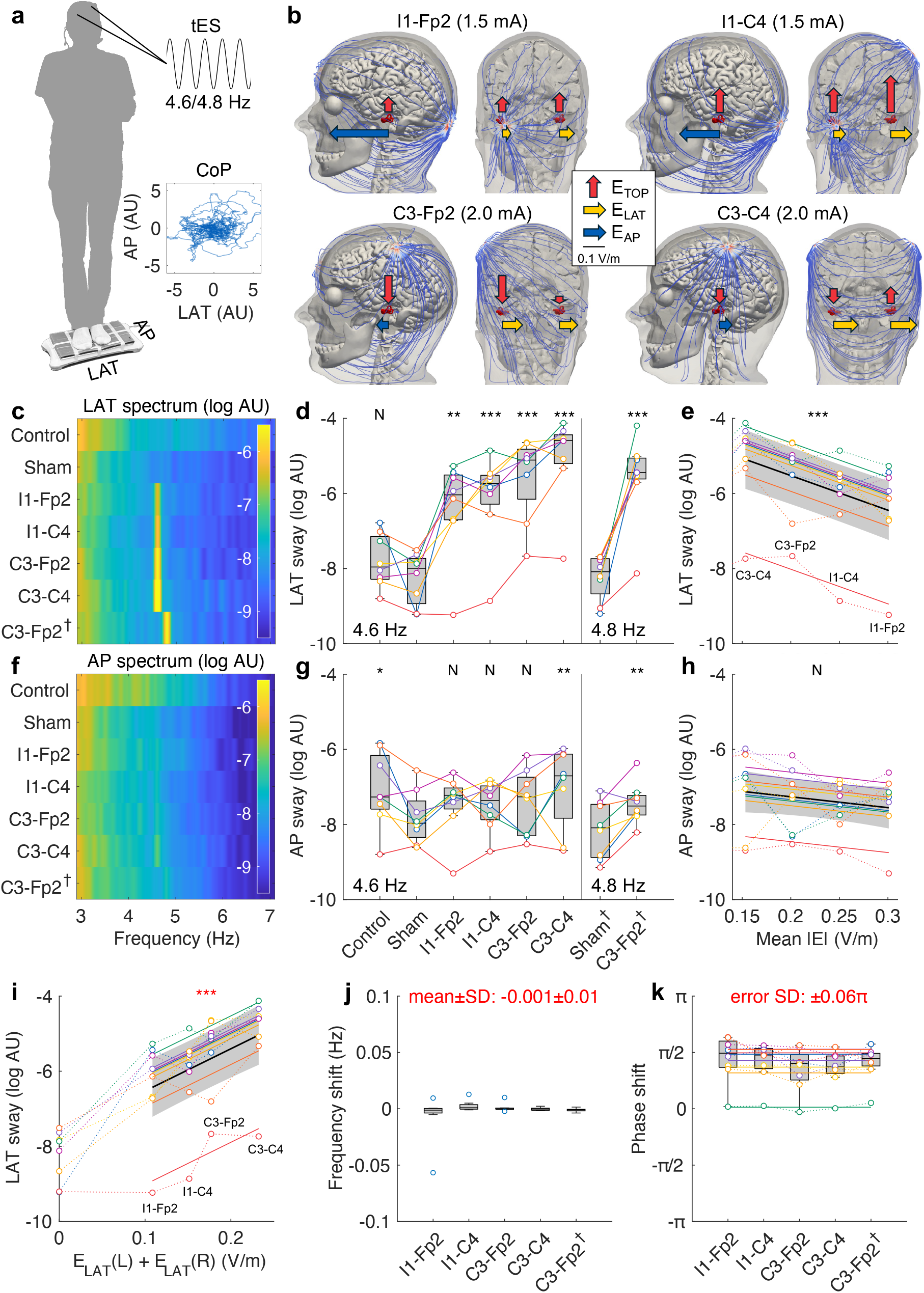
Body sway responses following tES with four electrode montages. **a**. The participants stood on a force platform, which measured the centre of pressure along the anterior-posterior (AP) and lateral (LAT) axes in arbitrary units (AU). The stimulus was a sine wave at 4.6 or 4.8 Hz. **b**. Modelled current streamlines for tES applied through four electrode montages, with the stimulation current intensity (peak) of each montage in parentheses. The arrows show the magnitude and relative direction of the electric field components in the left and right vestibular systems (mean values over 15 MRI-based models, Supplementary Fig. 1). **c**. Power spectra of the lateral sway signal averaged over eight participants shows peaks at the stimulation frequency. The stimulation frequency was 4.6 Hz for other active conditions and 4.8 Hz for the C3-Fp2^*†*^ condition. The control condition with no stimulation was unblinded and applied first in each experimental session. **d**. Lateral sway at the stimulation frequency increased compared with sham for all electrode montages (*n* = 8, paired t-test, control vs sham (two-sided): *t* = 1.50, *p* = 0.18, if excluding one outlier (blue): *t* = 1.26, *p* = 0.25; I1-Fp2, I1-C4, C3-Fp2, C3-C4, and C3-Fp2^*†*^ vs sham (one-sided): *t* = 4.68, *p* = 0.0011; *t* = 5.69, *p* = 0.00037; *t* = 6.92, *p* = 0.00011; *t* = 9.32, *p* = 1.7e-5; and *t* = 6.95, *p* = 0.00011). **e**. The average vestibular electric field magnitude was an invalid predictor for the lateral sway at 4.6 Hz (*n* = 8, likelihood ratio test, *χ*^2^(1) = 17.6, *p* = 2.7e-5). **f**. Average power spectra of the AP sway signal shows that the peak at the stimulation frequency is less visible than that for the lateral sway. **g**. The stimulation produced only small differences in the AP sway compared with sham (*n* = 8, paired t-test, control vs sham (two-sided): *t* = 2.48, *p* = 0.042; I1-Fp2, I1-C4, C3-Fp2, C3-C4, and C3-Fp2^*†*^ vs sham (one-sided): *t* = 1.70, *p* = 0.067; *t* = 0.95, *p* = 0.19; *t* = 1.62, *p* = 0.074; *t* = 3.47, *p* = 0.0052; and *t* = 3.68, *p* = 0.0039). **h**. There was no significant effect of average electric field magnitude on the AP sway at 4.6 Hz (*n* = 8, likelihood ratio test, *χ*^2^(1) = 2.35, *p* = 0.12). **i**. The lateral sway magnitude for the active conditions was proportional to the lateral electric field component, see yellow arrows in **b**. (*n* = 8, likelihood ratio test, *χ*^2^(1) = 19.5, *p* = 1.0e-5). **j**. The frequency of the lateral postural sway was equal to the stimulation frequency in all participants (*n* = 8, 95% CI: mean [−0.0068, 0.0009], SD [0.003, 0.0199]). **k**. The phase shift of the lateral sway compared with the stimulation waveform was almost constant across the active tES conditions within each participant (*n* = 8, 95% CI: SD [0.053*π*, 0.079*π*]). N *p >* 0.05, \**p <* 0.05, \*\**p <* 0.01, \*\*\**p <* 0.001. Black symbols indicate pre-registered analyses. Red symbols and text indicate exploratory analyses.

All electrode montages produced a significantly increased lateral body sway at the stimulation frequency compared with sham (Fig. 1d), with very large to huge effect sizes (Cohen’s *d*: 1.63, 1.94, 2.62, 3.00, and 2.61). The effect of stimulation on the AP sway (Fig. 1g) was smaller (Cohen’s *d*: 0.42–0.93) and, while some sway was visible in the spectrum (Fig. 1f), the effect might have been due to lateral sway, as not all participants stood exactly aligned with the axes of the force platform. No harmonics of the stimulation frequency were visible in the spectra (Fig. 1c,f and Supplementary Fig. 2).

We initially hypothesized that the sway magnitude would be proportional to the directionless electric field magnitude averaged over both vestibular systems. However, this hypothesis was clearly invalid, as the electrode montages with larger magnitudes produced a significantly smaller lateral sway (Fig. 1e). A potential explanation for the negative effect was that the vestibular systems were sensitive to the direction of the electric field in addition to its magnitude. Indeed, it appeared that the electrode montages with the largest sway were those with the largest lateral electric field component (Fig. 1b), which could be observed by fitting a linear mixed effect model (Fig. 1i). It must be noted that all bilateral sums or differences of the electric field components correlated strongly, either positively or negatively, among the four electrode montages (Supplementary Fig. 3a), and thus, any of them could be shown to have a statistically significant effect on the body sway. However, except for the sum of the bilateral lateral components (Fig. 1i), all other combinations could be easily shown to be invalid; either they would have a negative slope (Supplementary Fig. 3a), or predict a body sway for sham (Supplementary Fig. 3c,d).

Further exploratory analyses indicated that the peak of the sway signal precisely matched the stimulation frequency in all participants and conditions (Fig. 1j). Such precision would be very unlikely if higher-order physiological processes were involved, suggesting that the sway response was due to direct artificial stimulation, probably that of the vestibulospinal reflex. The phase of the sway with respect to the stimulus waveform was also consistent within each participant across the conditions (Fig. 1k), suggesting that the response originated from the stimulation of the same structure involving the same neural pathways for all conditions.

Subjective sensations reported by the participants in at least one condition were: skin sensations (8/8 participants), feeling swaying (4/8), feeling more stable or balanced (4/8), metallic taste (2/8), blinking lights (1/8), and rhythmic increase/decrease in background noise (1/8).

### Experiment 2

An unblinded additional experiment was conducted to investigate the body parts contributing to the body sway (Fig. 2a). Analysis of the horizontal sway from the video of three participants showed a lateral sway response in multiple body segments for both tES applied through the C3-Fp2 montage and EVS applied through the bilateral mastoid montage, with the largest response in the lower legs (Fig. 2b).

**Figure 2.**
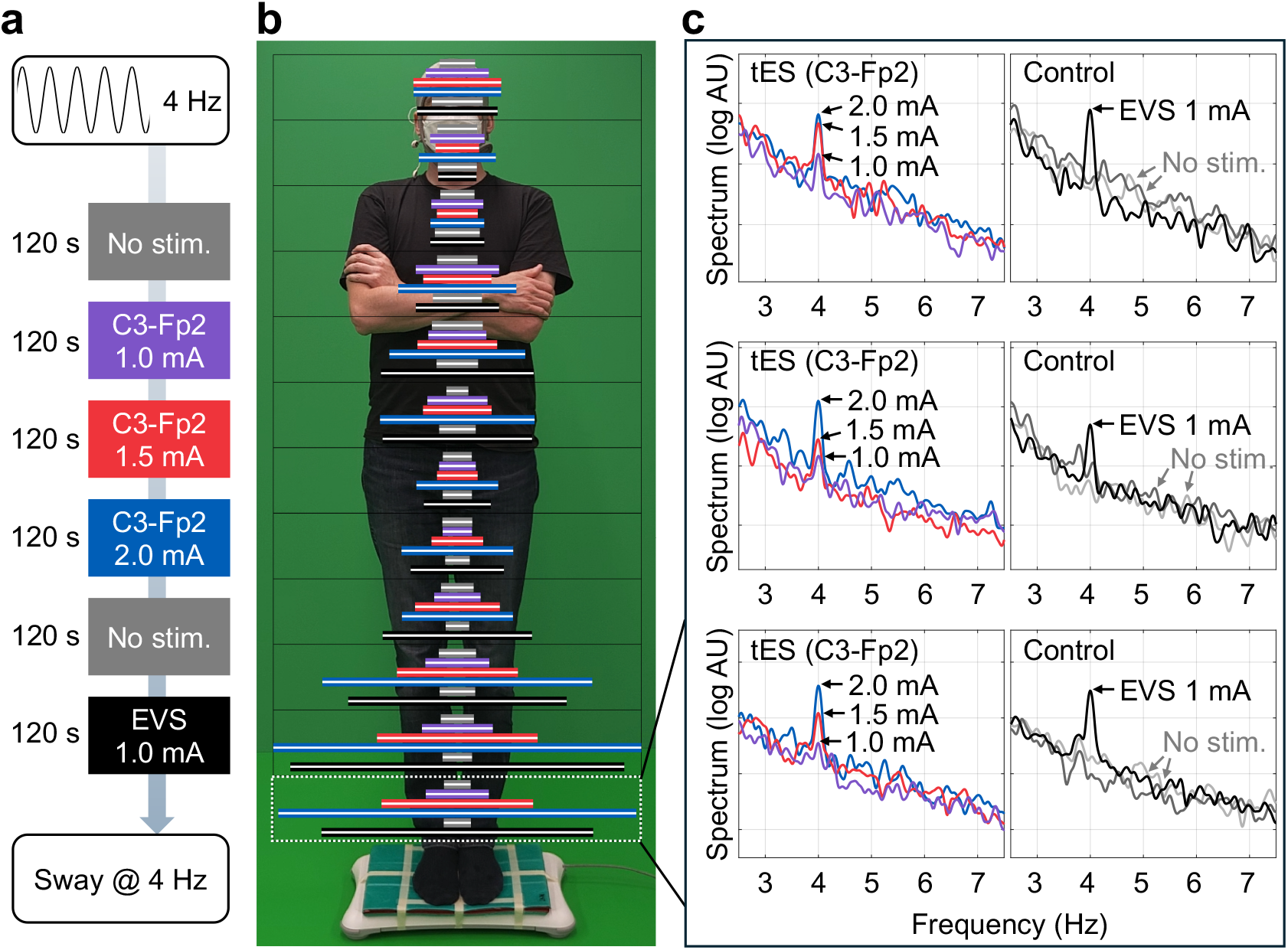
Video measurements of body sway responses for tES and EVS. **a**. Experimental conditions with a 4 Hz sine wave as the stimulus and the lateral body sway as the outcome. **b**. Horizontal bars indicate the magnitude of the 4 Hz spectral component (average over three participants) of the left-right sway signal in 12 body segments. The length of the bars is normalized, and their order and colour correspond to the conditions in **c**. One of the authors served as the model and gave consent for the publication of this picture in an online publication. **c**. Power spectra of the sway signal (arbitrary units, AU) from the lower legs of three participants (rows).

Similarly to experiment 1, the characteristic peak at the stimulation frequency, in this case 4 Hz, was present in all three participants (Fig. 2c). Current intensity of 1 mA applied to the C3-Fp2 montage was sufficient to produce a slight body sway at 4 Hz, and when the current intensity increased, the sway response increased, such that 2 mA produced roughly the same 4 Hz sway magnitude as 1 mA EVS. Modelling predicted that both montages produced approximately equal electric field magnitudes (Supplementary Fig. 1b) but that the EVS montage had 50% stronger lateral electric field component (Supplementary Fig. 1c). The distributions of the sway across body segments were similar for 2.0 mA tES with the C3-Fp2 montage and 1.0 mA EVS (Fig. 2b), suggesting a similar origin of the sway response for both conditions.

Participants reported subjective skin sensations (3/3) and metallic taste (1/3).

## Discussion

Alternating current applied to several commonly used tES electrode montages produced a body sway precisely at the stimulus frequency, with very large to huge effect size compared with sham. The sway magnitude was proportional to the calculated lateral component of the electric field within the vestibular system. The phase of the sway was locked to that of the stimulus, and the magnitude and distribution of the sway among the body segments was similar between C3-Fp2 and the bilateral mastoid montage, which is known to stimulate the vestibular system. The most likely explanation for these observations is that tES directly affected the vestibular system.

The effective frequency range of vestibular stimulation overlaps with common tES frequencies. The vestibular system is known to be sensitive to DC, leading to postural changes and sensations of illusory movements, observed since the 19th century [2, 1]. With AC, postural sway can be observed at least up to 10 Hz [3], and eye movements can be produced at least up to 20 Hz [9].

Therefore, we expect that the vestibular effects demonstrated here at 4–5 Hz are present in a wide frequency range covering both tDCS and theta-, alpha-, and lower beta-band tACS.

EVS activates a variety of upper limb, lower limb, and trunk muscles [1], which is also observable from our video posturography analysis showing increased sway both in the lower legs and central torso, which can potentially affect several outcomes related to motor function. The cerebral cortex is likely not involved in the immediate effects, as the earliest EVS responses are measured in lower leg muscles at latencies as short as 50 ms [10, 11]. It remains to be studied whether the vestibulospinal mechanisms causing muscle activations could interfere with the measurements of corticospinal excitability, e.g., the motor evoked potential (MEP) sizes measured using transcranial magnetic stimulation, which have been the main outcome measure in motor cortical tES studies [12]. Recently, MEP sizes have been shown to depend on the phase of 2.0 mA tACS at approximately 10 and 20 Hz [13], which could be due to vestibulospinal effects but also due to cortical excitability changes. Vestibular effects can also interfere with the measurement of tremor during tACS, which is typically measured using accelerometers [14, 15, 16] that can pick up postural sway responses. To robustly ensure that vestibular stimulation is not contributing to tES outcomes, EVS could be included as a positive control in experimental design.

The unintentional stimulation of the vestibular system is analogous to the sensation of flashing lights (phosphenes) observed during tACS [17], which has been attributed to the spread of the stimulation current to the retina [18, 19]. As with phosphenes, the unwanted vestibular effects can probably be reduced by designing electrode montages that cancel out the electric field in the vestibular system. Based on our exploratory analyses, the target for the minimization problem should be the lateral electric field component, as electrode montages with the smallest lateral component also minimized postural sway. However, the contribution of other electric field components cannot be ruled out, as they could have caused effects that were not visible as a postural sway. It is still an open question whether the electric field affects the vestibular hair cells, the vestibular afferent fibres, or both [2]. Notably, our macroscopic-level electric field modelling suggested that the field magnitudes were less than 0.3 V/m, which is at least an order-of-magnitude weaker than the field required to directly activate myelinated nerve fibres [20].

As far as the authors are aware, the body sway responses caused by tES have not been previously reported, which appears puzzling as the effect, e.g., for the commonly-used C3-Fp2 montage, was rather obvious in all participants. A likely reason is the context-sensitivity of EVS balance responses [1]. For example, the leg muscle responses observable in standing subjects disappear when the subjects are seated [1], which is the typical scenario in tES studies. In our measurement setup, we also attempted to maximize the signal-to-noise ratio of the sway response, performing the measurements in subjects standing eyes closed with their feet together, which has been shown to amplify leg muscle activity compared with a wider stance and open eyes [1, 21]. Furthermore, while the sway responses were easily observable with frequency-domain analysis of the force platform recordings or video, they were minute compared with normal body sway (*<* 2 Hz) and thus invisible to the naked eye and easily missed.

In conclusion, direct modulation of the vestibular system may interfere with the interpretation of tES studies. The effect is likely to exist for typical tES frequencies and current intensities as well as for multiple commonly used tES montages.

## Author contributions

Ilkka Laakso: Conceptualization, Methodology, Investigation, Formal analysis, Writing – Original Draft, Visualization, Supervision.

Janita Nissi: Methodology, Investigation, Formal analysis, Writing – Review & Editing, Visualization.

Otto Kangasmaa: Methodology, Investigation, Writing – Review & Editing.

---

**Supplementary Figure 1:**
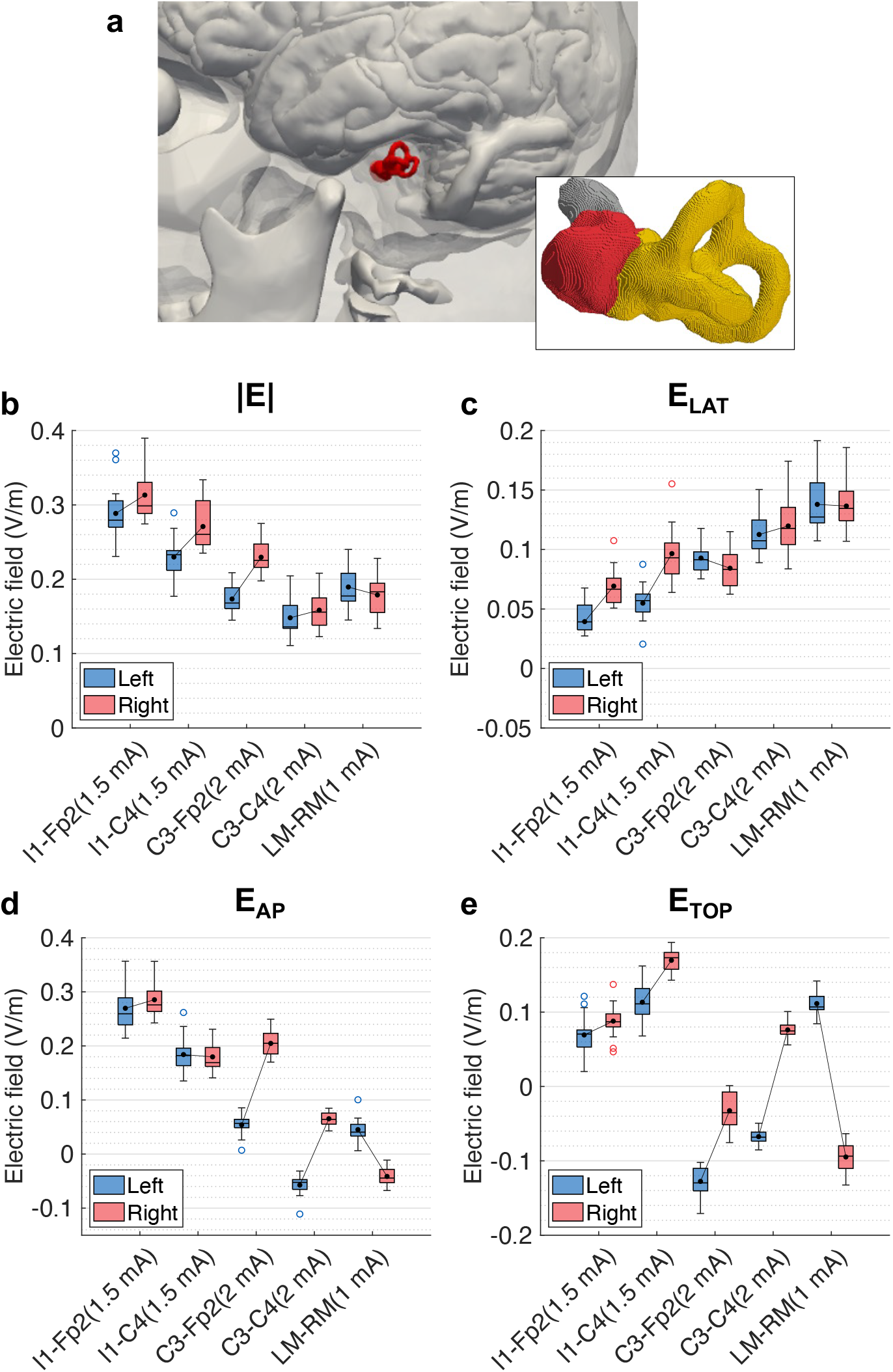
Calculated electric field data in 15 head models. **a**. Visualization of the inner ear model in a representative head model. The reported electric field values are arithmetic averages over the entire vestibular system model, shown in yellow. The vestibulo-cochlear nerve is shown in grey and the cochlea in red. (Illustration from: Nissi et al., J. Neural Eng. 21 (2024) 046038, doi: 10.1088/1741-2552/ad658f.) **b**. Box chart visualization of the electric field magnitude for five electrode montages used in this study, see Fig. 1b. LM-RM is the montage where the electrodes are placed over the bilateral mastoids, used in experiment 2. The mean values (black disks) of left and right vestibular systems are connected by a black line segment. **c, d, e**. Lateral, anterior–posterior, and superior–inferior electric field components at a positive peak of the stimulus sine wave. The positive directions are right, anterior, and superior, respectively. Note that a phase shift of *π* would flip all signs.

**Supplementary Figure 2:**
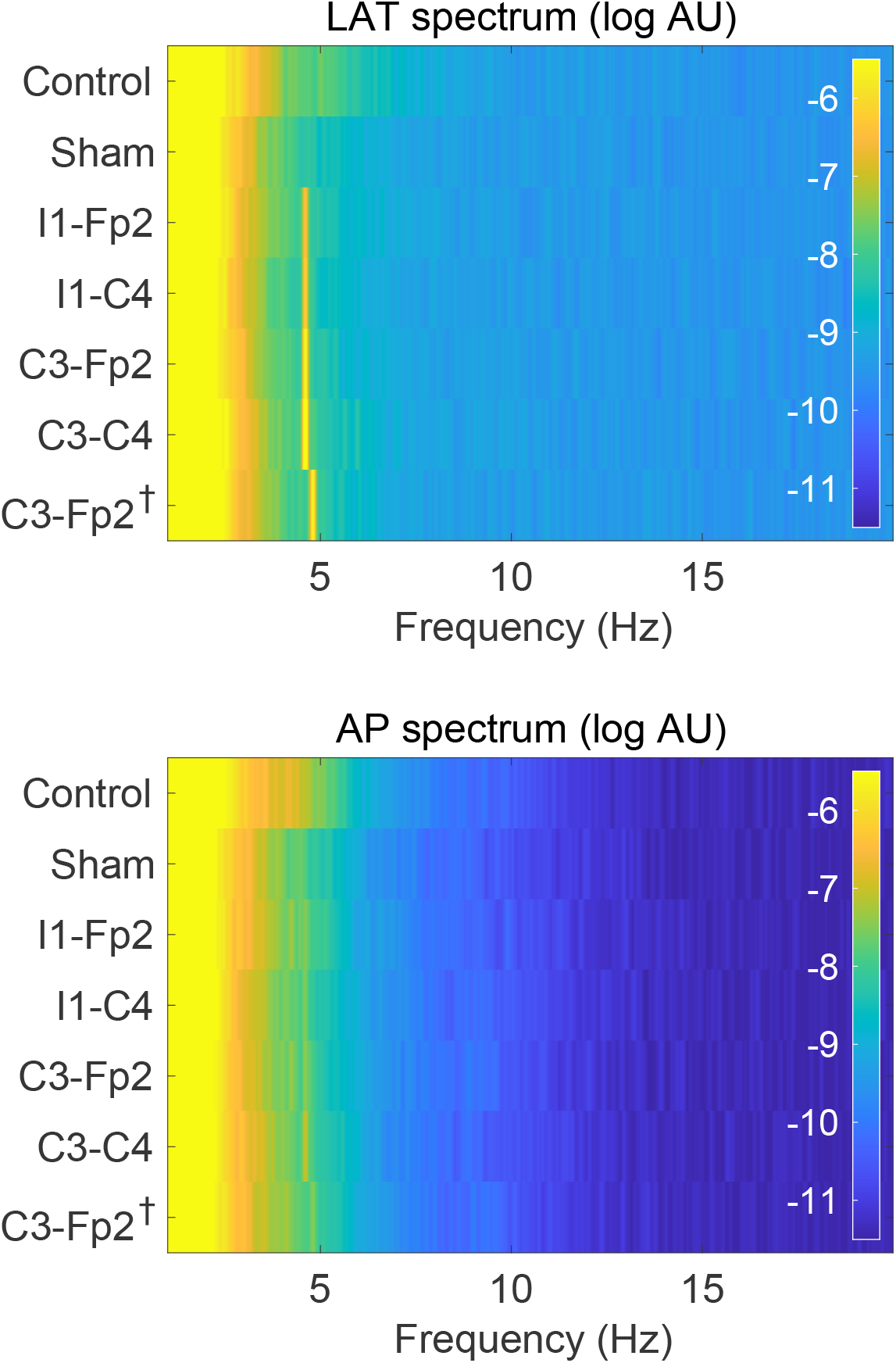
Average spectra of sway signals between 1 and 20 Hz. The data of Fig. 1c,f is visualized with wider frequency and amplitude ranges. Note that the colour scale is saturated at frequencies lower than approximately 3 Hz. Top: lateral sway, bottom: anterior-posterior sway.

**Supplementary Figure 3:**
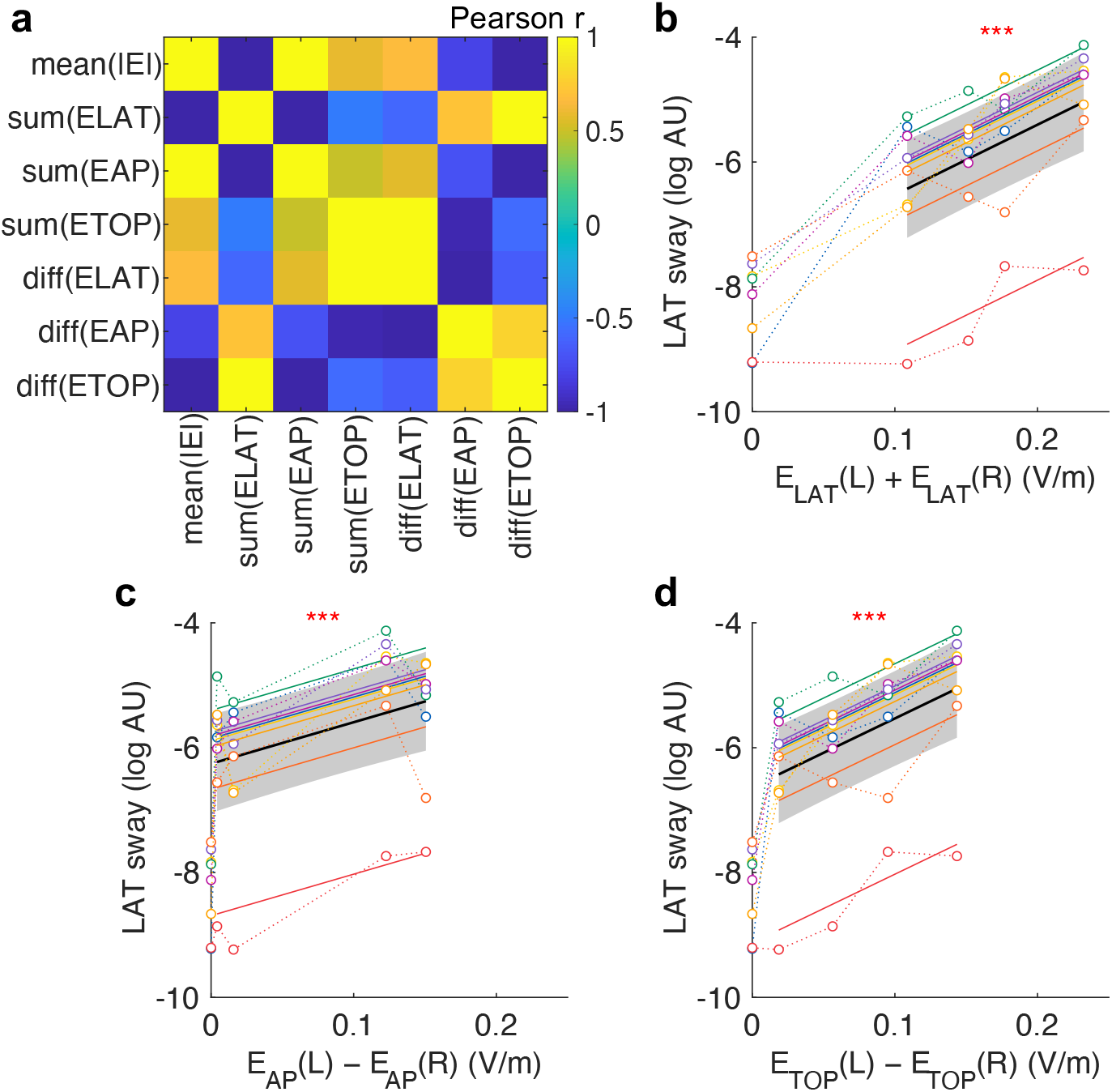
Exploratory analysis on the effect of electric field on the lateral body sway. **a**. Pairwise correlations of various electric field measures. The average magnitude and bilateral sums and differences of each electric field component were calculated. As the magnitude of the electric field correlated negatively with the lateral body sway (Fig. 1e), measures that correlated negatively with the magnitude had a positive effect on the body sway. All other measures are invalid, as they would predict a negative effect. **b, c, d**. Linear mixed effect models of dependence of the lateral sway on the sum of lateral electric field components (b), difference between the bilateral AP components (c), and difference between the bilateral superior-inferior components (d). In c and d, the models would predict a body sway for sham (electric field of 0 V/m), and thus, the models are invalid. Only the model with the lateral component (b) cannot be directly shown to be invalid. (*n* = 8, likelihood ratio tests, b: *χ*^2^(1) = 19.5, *p* = 1.0e-5, c: *χ*^2^(1) = 12.3, *p* = 0.00045, d: *χ*^2^(1) = 18.3, *p* = 1.9e-5.)

## References

[1] Fitzpatrick RC, Day BL. Probing the human vestibular system with galvanic stimulation. J. Appl. Physiol. 2004;96(6):2301–16.

[2] Dlugaiczyk J, Gensberger KD, Straka H. Galvanic vestibular stimulation: from basic concepts to clinical applications. J. Neurophysiol. 2019;121(6):2237–2255.

[3] Nissi J, Kangasmaa O, Kataja J, Bouisset N, Laakso I. In vivo and dosimetric investigation on electrical vestibular stimulation with frequency- and amplitude-modulated currents. J. Neural Eng. 2024;21(4):046038.

[4] Woods AJ, Antal A, Bikson M, Boggio PS, Brunoni AR, Celnik P, et al. A technical guide to tDCS, and related non-invasive brain stimulation tools. Clin. Neurophysiol. 2016;127:1031–1048.

[5] Oostenveld R, Praamstra P. The five percent electrode system for high-resolution eeg and erp measurements. Clin. Neurophysiol. 2001;112(4):713–719.

[6] Nissi J, Laakso I. Magneto- and electrophosphene thresholds in the retina: a dosimetry modeling study. Phys. Med. Biol. 2022;67(1):015001.

[7] Laakso I, Hirata A. Fast multigrid-based computation of the induced electric field for transcranial magnetic stimulation. Phys. Med. Biol. 2012;57(23):7753–65.

[8] Iacono MI, Neufeld E, Akinnagbe E, Bower K, Wolf J, Vogiatzis Oikonomidis I, et al. Mida: A multimodal imaging-based detailed anatomical model of the human head and neck. PLOS ONE 2015;10(4):e0124126.

[9] Mackenzie SW, Reynolds RF. Ocular torsion responses to sinusoidal electrical vestibular stimulation. J. Neurosci. Meth. 2018;294:116–121.

[10] Fitzpatrick R, Burke D, Gandevia SC. Task-dependent reflex responses and movement illusions evoked by galvanic vestibular stimulation in standing humans. J. Physiol. (Lond.) 1994;478(2):363–372.

[11] Rosengren S, Colebatch J. Differential effect of current rise time on short and medium latency vestibulospinal reflexes. Clin. Neurophysiol. 2002;113(8):1265–1272.

[12] Nitsche MA, Cohen LG, Wassermann EM, Priori A, Lang N, Antal A, et al. Transcranial direct current stimulation: State of the art 2008. Brain Stimul. 2008;1(3):206–23.

[13] Wischnewski M, Tran H, Zhao Z, Shirinpour S, Haigh ZJ, Rotteveel J, et al. Induced neural phase precession through exogenous electric fields. Nat. Commun. 2024;15(1).

[14] Brittain JS, Probert-Smith P, Aziz T, Brown P. Tremor suppression by rhythmic transcranial current stimulation. Curr. Biol. 2013;23(5):436–440.

[15] Mehta AR, Pogosyan A, Brown P, Brittain JS. Montage matters: the influence of transcranial alternating current stimulation on human physiological tremor. Brain Stimul. 2015;8(2):260–268.

[16] Schreglmann SR, Wang D, Peach RL, Li J, Zhang X, Latorre A, et al. Non-invasive suppression of essential tremor via phase-locked disruption of its temporal coherence. Nat. Commun. 2021;12(1).

[17] Kanai R, Chaieb L, Antal A, Walsh V, Paulus W. Frequency-dependent electrical stimulation of the visual cortex. Curr. Biol. 2008;18(23):1839–43.

[18] Kar K, Krekelberg B. Transcranial electrical stimulation over visual cortex evokes phosphenes with a retinal origin. J. Neurophysiol. 2012;108(8):2173–8.

[19] Laakso I, Hirata A. Computational analysis shows why transcranial alternating current stimulation induces retinal phosphenes. J. Neural Eng. 2013;10(4):046009.

[20] Reilly JP. Peripheral nerve stimulation by induced electric currents: Exposure to time-varying magnetic fields. Med. Biol. Eng. Comput. 1989;27(2):101–110.

[21] Day BL, Steiger MJ, Thompson PD, Marsden CD. Effect of vision and stance width on human body motion when standing: implications for afferent control of lateral sway. J. Physiol. (Lond.) 1993;469(1):479–499.

